# Exploring the limits of learning: segregation of information integration and response selection is required for learning a serial reversal task

**DOI:** 10.1101/163725

**Authors:** Camilo J. Mininni, B. Silvano Zanutto

## Abstract

Animals are proposed to learn the latent rules governing their environment in order to maximize their chances of survival. However, rules may change without notice, forcing animals to keep a memory of which one is currently at work. Rule switching can lead to situations in which the same stimulus/response pairing is positively and negatively rewarded in the long run, depending on variables that are not accessible to the animal. This fact rises questions on how neural systems are capable of reinforcement learning in environments where the reinforcement is inconsistent. Here we address this issue by asking about which aspects of connectivity, neural excitability and synaptic plasticity are key for a very general, stochastic spiking neural network model to solve a task in which rules change without being cued, taking the serial reversal task (SRT) as paradigm. Contrary to what could be expected, we found strong limitations for biologically plausible networks to solve the SRT. Especially, we proved that no network of neurons can learn a SRT if it is a single neural population that integrates stimuli information and at the same time is responsible of choosing the behavioural response. This limitation is independent of the number of neurons, neuronal dynamics or plasticity rules, and arises from the fact that plasticity is locally computed at each synapse, and that synaptic changes and neuronal activity are mutually dependent processes. We propose and characterize a spiking neural network model that solves the SRT, which relies on separating the functions of stimuli integration and response selection. The model suggests that experimental efforts to understand neural function should focus on the characterization of neural circuits according to their connectivity, neural dynamics, and the degree of modulation of synaptic plasticity with reward.

## Introduction

Natural environments are complex places in which animals strive to survive, with hidden variables and stochastic factors such that the information available at any moment is partial, and it must be sampled at several time points and integrated. What is more, the rules governing the environment might change with time, leading to conflicting information. For example, an animal might learn how and where to seek for food, but if the place for feeding cyclically changes, or the means of obtaining food change, the animal has to switch strategies along [1]. In this case, no unique strategies exist, but several strategies must be learned. More importantly, the value of a response not only depends on the current scenario, but in the history of events, for example, the history of recent success of a given strategy. Therefore, it is relevant to study tasks in which rules might change over time and thus reinforcement of stimulus/response pairings might be inconsistent, i.e. Inconsistent-Reinforcement tasks (IRTs). Learning an IRT through a neural network model can be problematic: since each stimulus/response pairing is positively and negatively reinforced in the long run, learning of one rule may lead to the erasure of information regarding other rules, conforming a case of catastrophic forgetting [2]. Thus, it is of special interest to understand the neural mechanisms involved during learning of this kind of tasks.

The goal of this work is to find the essential properties required by biologically plausible neural networks to solve a Serial Reversal Task (SRT), which is an IRT where two rules alternate over time, demanding the animal to keep track of previous events in order to maximize reward. We focus on stochastic spiking neural networks (SSNN), a very general kind of neural network model that has been employed to explain how key features of neural circuits, like excitatory-inhibitory balance [3] and spike timing-dependent plasticity (STDP) [4], can lead to Bayesian inference [5] and reinforcement learning [6]. For a very general family of SSNNs, we show analytically that strong limitations to learning the SRT emerges when the functions of integration of stimuli information and response selection are conducted by the same neural population. We propose a model that is able to learn the SRT and discuss the implications of the results regarding the neural mechanisms of decision-making.

## Results

We will study the characteristics of an agent controlled by a biologically plausible neural network that learns to solve a SRT, conforming to what we will define as the hypothesis of functionality by learning, which states that the set of configurations that gives functionality is a small subset of the set of initial configurations. In this way, functionality is acquired by a learning mechanisms that always leads the system from any random initial condition to one of the functional configurations. The hypothesis implies that the system is not initially designed to solve a given task from start.

A SRT is a discrimination task in which the mapping between the stimulus and the correct response is reversed after a given (random) number of trials (Fig. 1a). One out of two possible cue stimuli (*s*
_1_ or *s*
_2_) is presented to the agent. During cue presentation the agent has to execute one out of two possible responses (*R*
_1_ or *R*
_2_) in order to get a reward. Which response is correct depends on the current rule (rule *L*
_1_: *s*
_1_ → *R*
_1_, *s*
_2_ → *R*
_2_; rule *L*
_2_: *s*
_1_ → *R*
_2_, *s*
_2_ → *R*
_1_). A reward stimulus is shown after cue presentation: *r*
_1_ for correct responses or *r*
_0_ for incorrect ones. One rule withstand until a switch of rules occurs at random. Switching occurs with low probability, to ensure that a considerable number of trials with the same rule are presented.

**Fig. 1.**
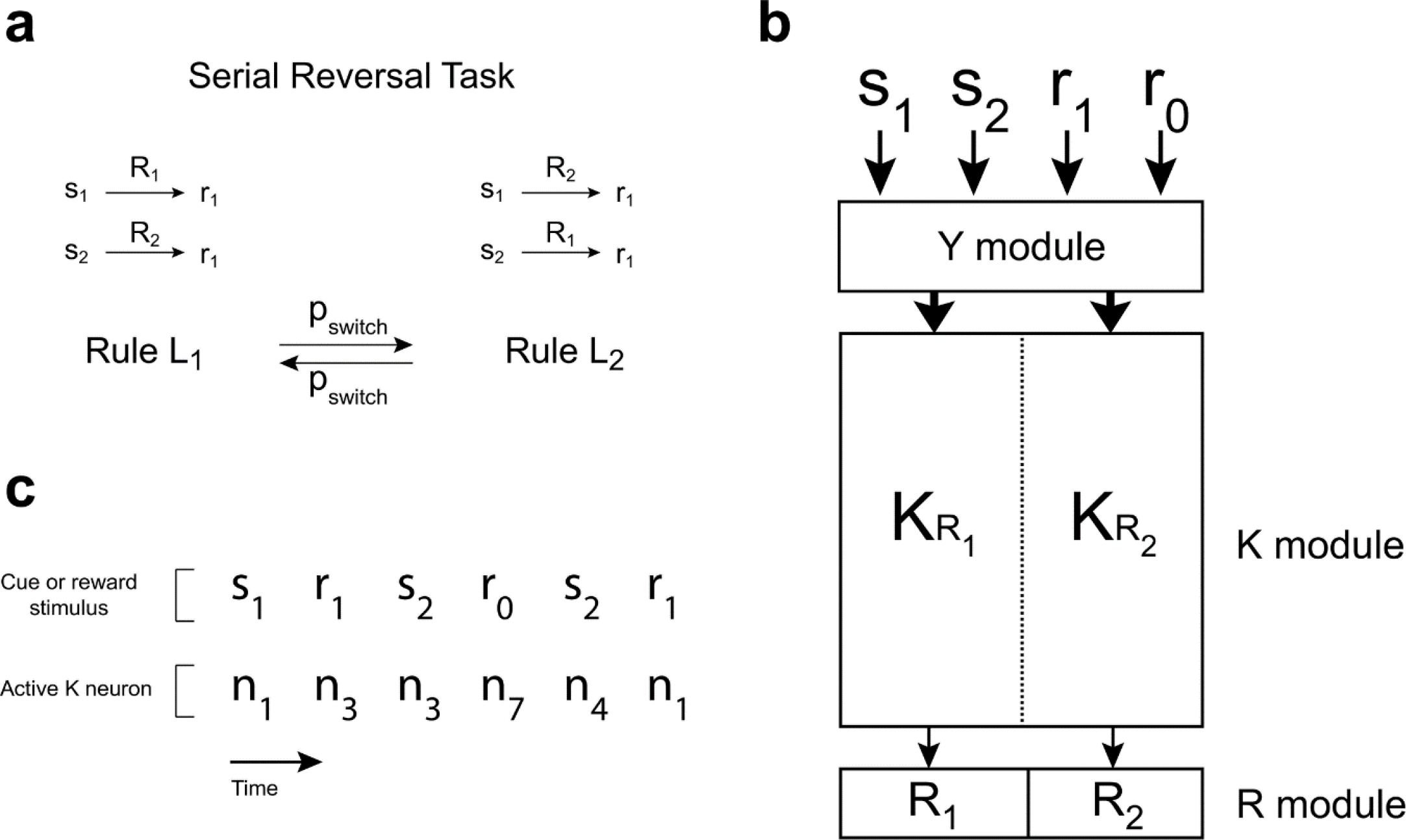
Serial Reversal protocol and simple network connectivity. (**a)** Each trial is composed of a cue stimulus presentation, during which the behavioural response must be executed, and a reward stimulus presentation. Correct responses depend on the stimulus presented and the current rule, which changes with probability *p switch*. (**b)** Diagram representing the general connectivity of the simple network. Neurons in module *Y* codifies both cue and reward stimuli, and projects to the *K* module. *K* neurons connect with each other and projects to one of the two response neurons. Therefore, *K _R_1_
_
* neurons can be sorted in two halves depending on whether they project to *R*
_1_ neuron (*K* neurons) or *R*
_2_ neuron (*K_R_2_
_
* neurons). Firing of any *K* neuron elicits their target *R* neuron to fire. Connections between module *K* and module *R* are assumed to be hardwired prior to any learning, such that firing of *K* neurons completely defines the executed response. (**c)** An example sequence of 3 trials of a SRT for the model depicted in (b) prior to learning, with a minimal *K* module composed of 8 neurons.

**Fig. 2.**
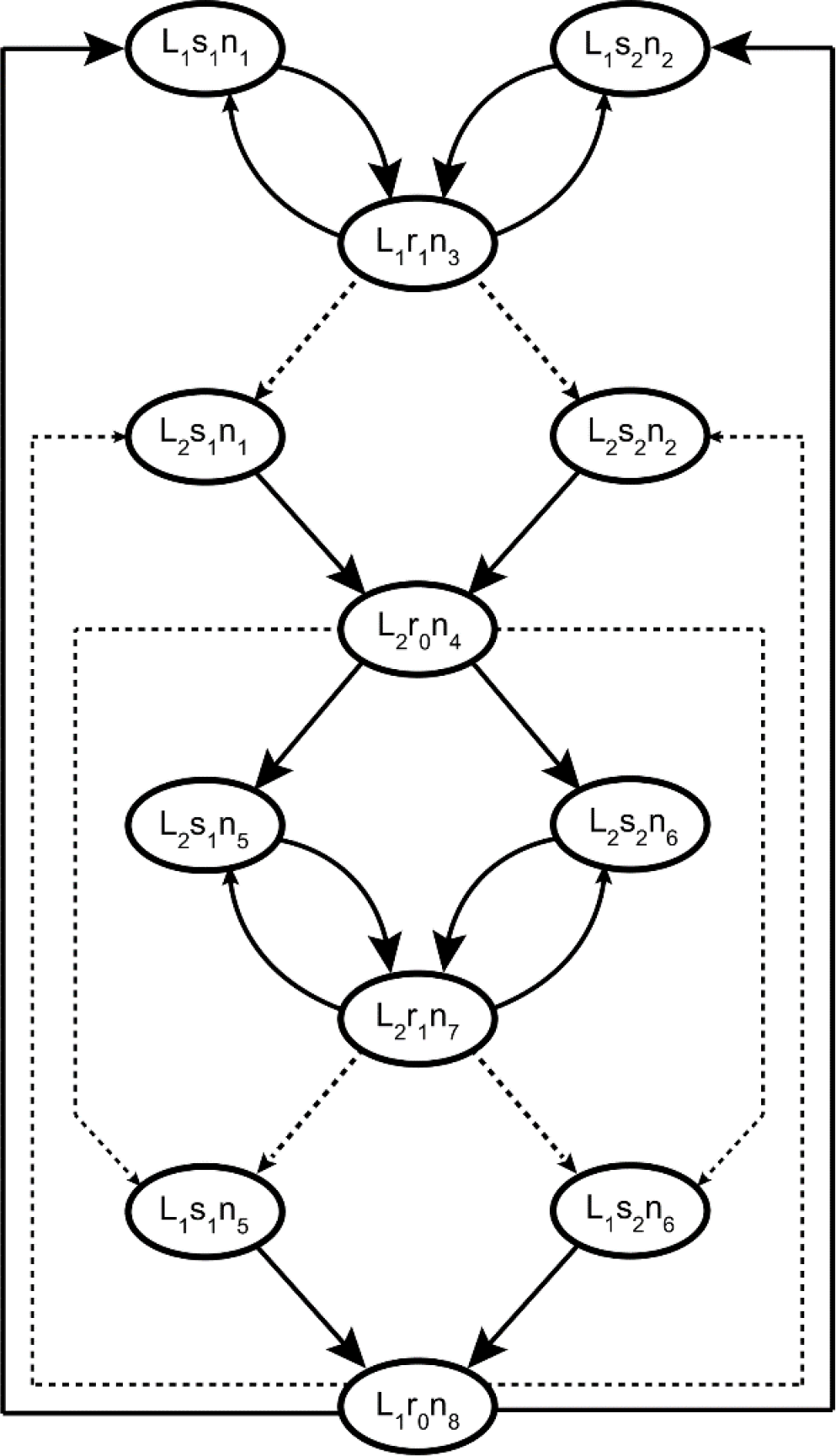
Graph representing the transition probabilities of the Markov chain associated with the simple network of Fig. 1 solving the SRT. The active *Y* neuron and the *R* neuron are excluded from the global state to simplify the representation, since *Y* neurons are entirely defined by the stimulus, and the *R* neurons are entirely defined by the active *K* neuron. The size of the arrow head represents the magnitude of the transition probability. Dashed lines depict transitions for which a change of rule occurs. Possible transitions that have no arrow are considered to have very low probability. The model has one set of stimuli coding neurons for each rule, and one neuron per rule to elicit the transition between rules when an error occurs.

The structure of the task implies that any agent that follows only one stimulus/response mapping as strategy will fail to get reward in half of the trials. Moreover, information provided by the stimuli is useless unless the agent is capable of retaining information about the current rule. Optimal performance can be achieved by adhering to a successful strategy, and to switch strategies when the current one is no longer successful.

We will consider an agent that is controlled by a spiking stochastic neural network composed of a sensory module *Y* and an integration/decision module *K* (Fig. 1b). Neurons in module *Y* code the sensory stimuli and project to module *K*, while neurons in module *K* project to the response neurons and to other *K* neurons. One half of the *K* population projects to response neuron *R* (the *K* _
*R*
_1_
_ subset of module *K*), the other half to response neuron *R*
_2_ (the *K* _
*R*
_2_
_ subset of module *K*). We assume that the firing of any neuron within a *K_R_
* group is enough to trigger the corresponding behavioural response. Therefore, the *K* module integrates sensory information together with information from within the network, and at the same time it defines the response that is going to be executed.

We assume that the neural network sketched in Fig. 1b fulfils the Markov condition: the probability of transitioning to a given state is only defined by the current state. This means that information about past events can only by carried on in the current state of the system. In the case of a SRT, a same stimulus should elicit either the responses *R*
_1_ or *R*
_2_, depending on the current rule. For example, *s*
_1_ should elicit response *R*
_1_ only during rule *L*
_1_, or *R*
_2_ only during rule *L*
_2_. This implies that *s* should elicit a response from a subset of the *K* group when *L*_1_ rule is current, or from the *K* group when *L*
_2_ rule is current. Since there is no explicit stimulus acting as a cue of the rule, the differential response of the *K* module in front of the same stimulus can be achieved only if the *K* neurons integrate inputs from the *Y* module together with inputs from the *K* module itself. This means that each stimuli must be coded by different groups of *K* neurons depending on the current rule. Then, the occurrence of an error should act as pivot, leading the system to the set of states associated with the other strategy.

In what follows we will show that the network sketched in Fig. 1b (referred to as *simple network*) is incapable of learning to solve the SRT without contradicting the hypothesis of functionality by learning. First we will consider a “reduced” example of the simple network of Fig. 1b that nevertheless puts in evidence the nature of the problem (see *Methods* section for a proof regarding both the reduced network and a general version of the simple network).

The firing state of module *K* will be represented by a vector *n*(*t*), where each element *n_i_
* (*t*) ∈{0,1} represent the firing state of the ith *K* neuron. Indistinctly *n_i_
* (*t*) will be used to indicate that the ith *K* neuron is active. Similarly, we define a vector *y*(*t*) where each element *y_i_
* (*t*) ∈{0,1} represents the state of neuron *y_i_
*; *y_i_
* will also be used to indicate that neuron *y_i_
* is active.

We will consider a network with a *Y* module composed of 4 neurons such that each stimulus is perfectly codified by one specific neuron, i.e. (*p*(*y_i_
* | *S_i_
*) = 1 and *p*(*y_i_
* | *S _j_
*) = 0 ∀*i* ≠ *j*) where *S_i_
* is the ith element of *S* = (*s*
_1_, *s*
_2_, *r*
_1_, *r*
_0_). Module *K* is composed of 8 neurons, which is the minimum number of neurons required to solve the SRT: one neuron for each stimulus (cue or reward) for each rule. Each trial *T* has two time points (*t* and *t* +1), one for cue presentation and another for reward stimulus presentation. The *K* _
*R*
_1_
_ group comprises neurons from 1 to 4; *K* _
*R*
_2_
_ comprises neurons from 5 to 8. At each time point only one *Y* neuron and one *K* neuron fires, and the decision is evaluated during cue presentation (Fig. 1c). Then, each neuron in module *K* has a probability of firing that is given by:

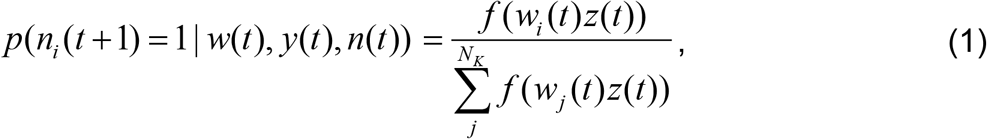

where *w* stands for all synaptic weights in the network, *w_i_
* is a vector containing the synaptic weights of afferent connections from all *Y* and *K* neurons onto the ith neuron in module *K*, and *z* is vector containing the firing states of all *Y* and *K* neurons such that *w_ij_
* is the synaptic weight of the jth neuron with firing state *z _j_
* that projects to neuron *i*. The function *f* can be any function with the sole condition of being strictly increasing with *w_ij_
*. Equation (1) endorses the *K* module with characteristics of a “soft winner-take-all” circuit in which a highly excited neuron inhibits the other neurons in the module through a global inhibitory circuit [5].

Synaptic weights *w_ij_
* change according to the local pre/post synaptic activity and the reward stimuli *r*. The change Δ*w_ij_
* of a synaptic weight *w_ij_
* is given by a function *g*:

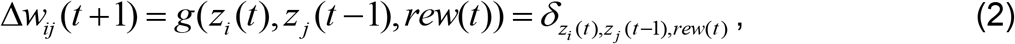

Where *z_i_
* (*t*) and *z _j_
* (*t* −1) are the respective pre and post synaptic states, and *rew* is a function of the delivery of reward, such that *rew* = 1 during the cue and reward presentation for trials in which the response was correct, and *rew* = 0 otherwise. The *g* function can be in principle any function adopting real values *δ*.

Now we can write the transition probability of the Markov chain that describes the dynamic of the whole system (network, stimuli and rules):

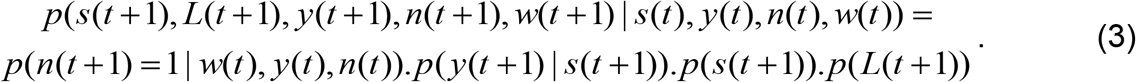

Equation (3) is obtained by applying the chain rule of conditional probabilities, and using the fact that *L* is independent of stimulus, and that firing state *n*(*t* +1) is independent of any other variable when conditioned to *n*(*t*), *y*(*t*) and *w*(*t*). Note that, since plasticity is assumed deterministic, eq. (3) is true if *w*(*t* +1) is the resulting synaptic weight configuration of applying function *g* given (*n*(*t*), *y*(*t*), *n*(*t* +1)). Any transition to different synaptic weight configuration will have zero probability.

Now we can find the transition probabilities that solve the SRT and study under what conditions a learning process is capable of reaching the solution. Figure 2 shows the directed graph for the transitions in the state space that solve the SRT. Under rule *L*
_1_ neurons *n*
_1_ and *n*
_2_ fire with cue *s*
_1_ and cue *s*
_2_ respectively, while neuron *n*
_3_ codes *r*
_1_ and *n*
_4_ codes *r*
_0_. For rule *L*
_2_, neurons *n*
_5_ and *n*
_6_ fire with cue *s*
_1_ and cue *s*
_2_, while neuron *n*
_7_ codes *r*
_1_ and *n*
_8_ codes *r*
_0_. Neurons *n*
_4_ and *n*
_8_ are responsible for the strategy switching in the behaviour of the agent. Each time a transition between rules occurs, an error is committed, and the corresponding error neuron fires. Equation 1 tells us that the only way to change the transition probabilities is by adjusting the synaptic weights. Since the *f* function is strictly increasing with *w_ij_
*, weights must be increased to favour a transition, or decreased to make a transition less probable. For example, the synaptic weights configuration must ensure that only neuron *n*
_3_ fires when (*r*
_1_, *n*
_1_) or (*r*
_1_, *n*
_2_) are the presynaptic active neurons (Fig. 3).

**Fig. 3.**
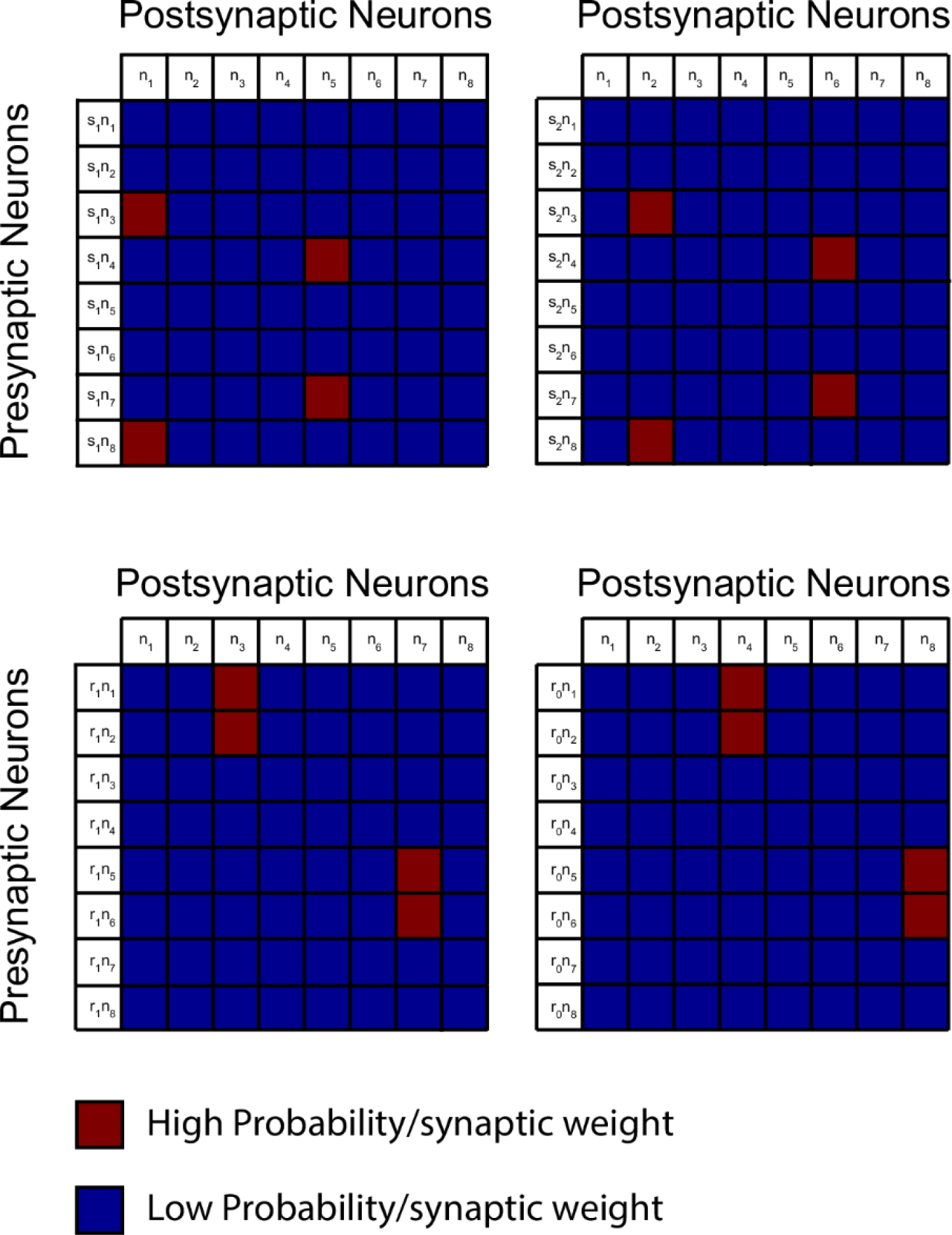
Specificity of module *K* responses given the presynaptic neurons firing. The matrixes show the probabilities/synaptic weights of postsynaptic *K* neurons being active according to the state of the presynaptic *K* and *Y* neurons that where active in the previous time step. Probability magnitudes are consistent with the Markov chain of Fig. 2. This representation gives a hint about how the synaptic weights ought to be. High transition probabilities can be achieved by setting high synaptic weights between a given presynaptic pair and the target postsynaptic neuron, and low synaptic weights for all other postsynaptic neurons.

It is evident that the transition matrix that solves the SRT has very specific probabilities, in which some transitions must not occur if other transition does. Conversely, this specificity is translated to the synaptic weight matrix that is solution for the SRT (Fig. 4a). The network will learn if the plasticity function *g* leads the system to the solution weight matrix regardless of the initial conditions, something that is questionable given that for the SRT any stimulus response pairing can be rewarded. To understand this, Fig. 4b shows the solution weight matrix for a discrimination task (DT) with two rules like in the SRT, but with the difference that rules are cued, so that the *Y* module can codify them. In this case, the set of stimulus/response pairings that leads to reward and the set that leads to no reward are disjoint sets. This fact is what makes possible to find a network that converge to the solution matrix for this DT by choosing a suitable *g* function (for example a Hebbian plasticity rule) that leads to increments in the synaptic weights only when a reward is obtained. But in the case of the SRT there are no disjoint sets of stimulus response pairings that separates reward from no reward. In fact, since we assume that the system is initiated without any information about how to solve the task, it can be seen that: 
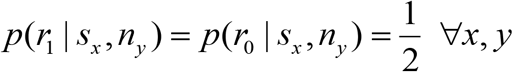
 and *p(n|r_1_) = p(n|r_0_) = p(n)* In particular:

**Fig. 4.**
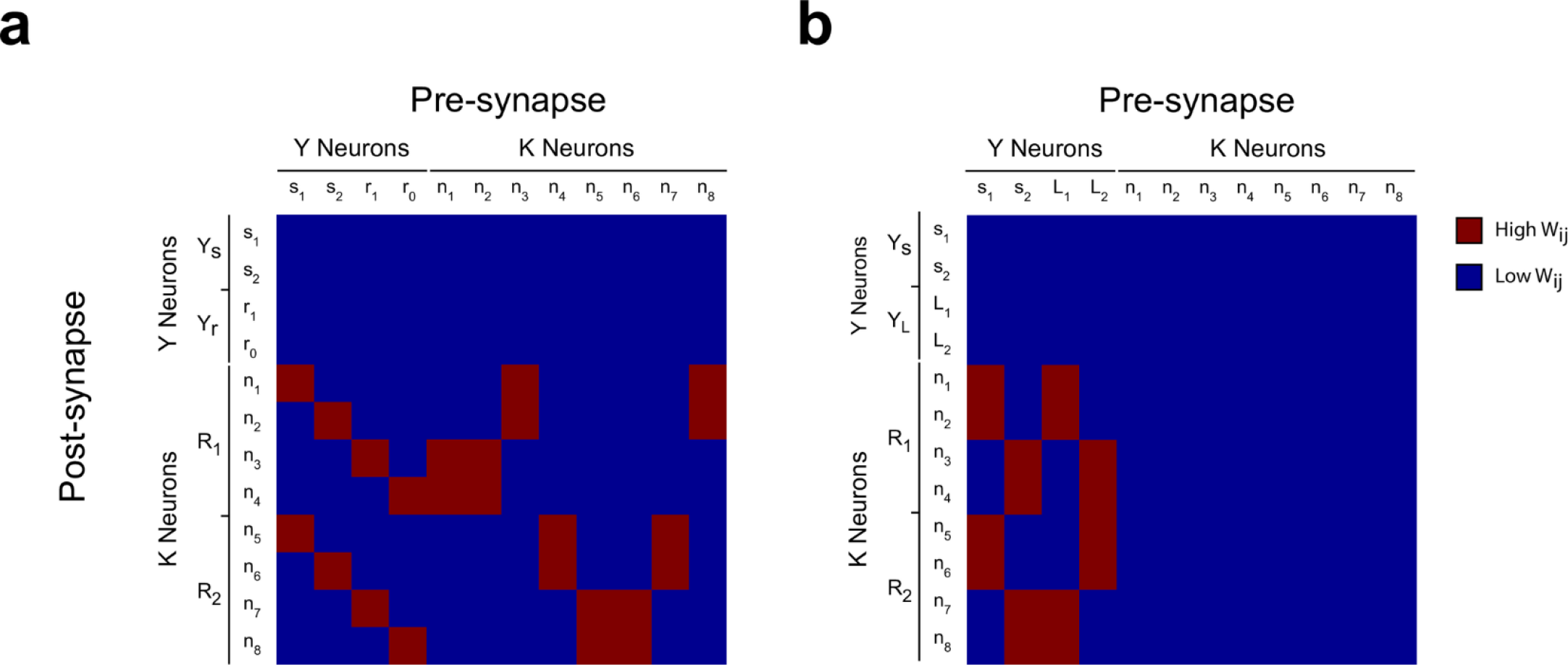
Synaptic weights between neurons of *Y* and *K* modules. (**a)** Synaptic weight configuration that allows the model to solve the SRT, consistent with the transition probabilities shown in Fig. 2. It can be seen that a specific arrangement of synaptic weights are required. (**b)** Synaptic weight configuration that allows the model to solve a DT. In contrast with the SRT, all high synaptic weights correspond to pre-post synaptic neurons that are systematically active when reward is obtained.

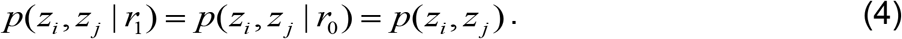

This allows us to write the average change 〈Δ*w_ij_
* 〉 for a given *w_ij_
*:

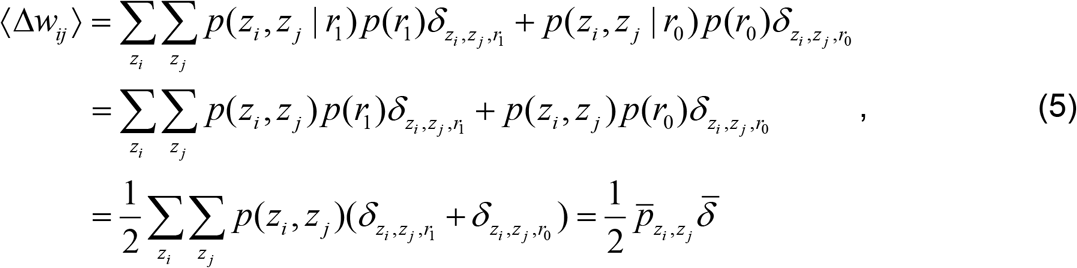

where 
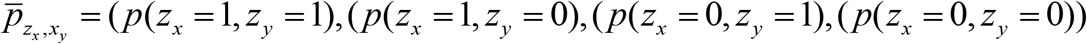
 and δ = (*δ*
_111_ +*δ*
_110_, *δ*
_101_ +*δ*
_100_, *δ*
_011_ +*δ*
_010_, *δ*
_001_ +*δ*
_000_).

As expected, 〈Δ*w_ij_
* 〉 is independent of the occurrence of reward. The 〈Δ*w_ij_
* 〉 can be understood as the inner product between the vector 
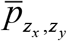
 representing the probability distribution of the pre/post synapsis pair, and *
δ
*, which contains the net change in *w_ij_
* for each pre/post configuration. The inner product implies a kind of correlation between the two vectors, and changing two *w_ij_
*: in specific directions requires specific adjustment of this inner product

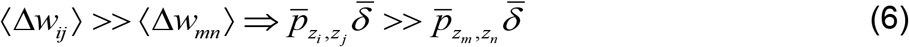

Thus, to get to the solution weight matrix pictured in Fig. 4a, a detailed adjustment between the probability distribution of *n*(*t*) and the plasticity function must hold. Adjusting *
δ
* to 
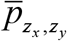
 would mean that the plasticity was designed to solve a specific task for a specific initial condition, contradicting the hypothesis of functionality by learning. Adjusting 
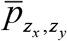
 to *
δ
* would mean that the initial synaptic weights were specifically chosen to solve the task, again contradicting the hypothesis. Therefore, since the requirements for reaching the *w_ij_
* that are solution to the SRT necessarily contradicts the hypothesis, we must conclude that the neural network sketched in Fig.1b and described by equations 1,2 and 3 cannot learn to solve a SRT.

### Learning to solve a SRT requires segregation of stimulus history coding from decision making

The incapacity of the model depicted in Fig. 1b for solving the SRT stems from the fact that the solution weight matrix cannot be reached by any plasticity function *g*. Conversely, this characteristic arises from two facts:
1. Correct stimulus/response pairings change over time, and there are no cues that give information about the current rule. Thus, in order to keep information about the current rule, the responsivity of the system towards the stimuli must be specially conditioned by the previous states of the system.
2. The population that codes information about the current rule is the same population that defines the behavioural motor response.

Fact number 1 implies that the task cannot be solved as a DT, since the reward does not separate stimulus/response pairs into any two disjoint subsets. Fact number 2 implies that coding of stimuli cannot be done freely, because when a neuron codes a stimulus by firing, it is also defining a motor response that is expected to lead to reward. Fact number 1 cannot be skipped because it originates from the very nature of the task. But fact number 2 can be circumvented in a model in which coding and decision functions are performed in separated neural populations. Figure 5a depicts such a model (referred to as *complex network;* see Methods for description of the implementation). There, module *K* integrates information about cues and reward as before, and about the response executed as well, but does not defines the motor response. Neurons in the integration module *K* project to two decision neurons *D*
_1_ and *D*
_2_. Which decision neuron fires univocally defines which response neuron (*R*
_1_ or *R*
_2_) will activate, leading to the corresponding motor response.

Therefore, module *K* needs to codify all the information required to solve the task. Ideally, it would suffice that neurons in module *K* codified the cue presented and the current rule. Nevertheless, no cue informs about the current rule, and module *K* only sees stimuli. Therefore, information about the current rule must be extracted from the history of perceived stimuli. For example, the sequence (*s*
_1_, *R*
_1_, *r*
_1_) shows that rule *L*
_1_ was currently working, and it should continue to do so except a reversal occurs, which is unpredictable but relative rare. In this manner, a possible solution is that neurons in module *K* codify each stimulus differently, depending on the previous stimulus history or contingency. This can be done following the model presented in Kappel *et al* [7]. There, it was shown that spike timing-dependent plasticity (STDP) in a soft winner-take-all stochastic neural network leads to the formation of groups of neurons that code stimuli distinctively, depending on stimuli history. In our case, module *K* should divide in groups of neurons codifying sequences of 4 stimuli: (*s*(*T* −1), *R*(*T* −1), *r*(*T* −1), *s*(*T*)), implying 16 possible contingencies.

The SRT structure for the following simulations is depicted in Fig. 5b. Each trial starts with the presentation of one cue, 25 ms long. At *t_decision_
* = 15 ms from trial onset, the state of neurons in the *R* module are actualized based on which neuron is firing in module *D*. At the same time, the response is characterized as correct or incorrect. During the interval [15 ms,25 ms] the synapsis from module *K* to module *D* are modified following eq. (24). The state of the *R* neurons are sustained unaltered between actualizations. The reward stimulus, also 25 ms long, is presented immediately after cue offset, being *r*
_1_ or *r*
_0_ depending on the correctness of the response. The rule is reversed every 15-20 trials, unless otherwise stated.

**Fig. 5.**
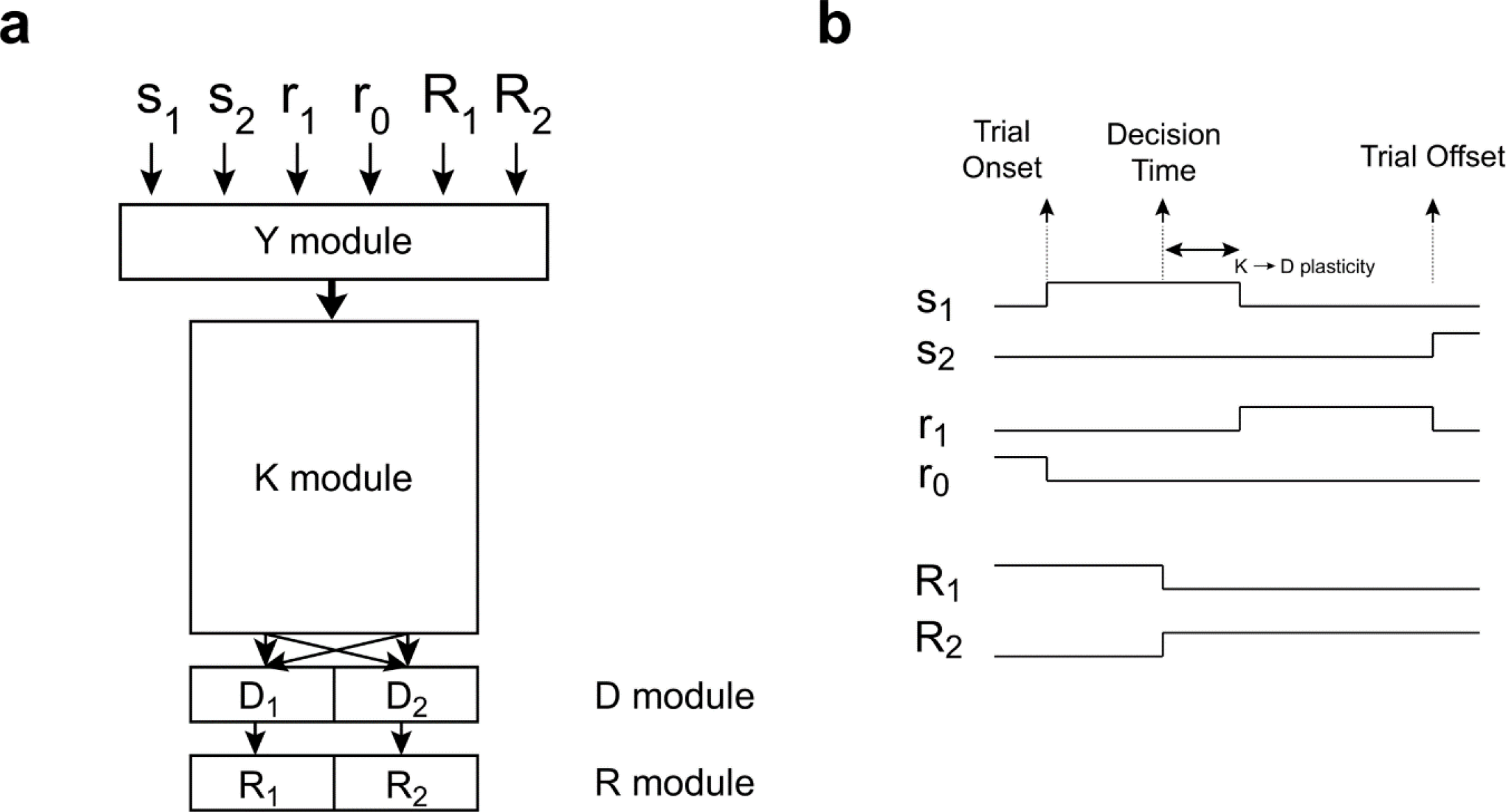
Serial Reversal protocol and complex network connectivity. (**a)** Diagram representing the general connectivity of the complex network. Each cue and reward stimulus is coded by the *Y* neuron population, like in the simple network. Besides, the executed motor response gives sensory feedback, such that each response is also coded by module *Y*. Module *Y* connects to all neurons in the integration module *K*, which in turn connect with each other and with each neuron in the decision module *D*. Each neuron *D* is hardwired to one neuron *R*, so that the response executed is entirely defined by the *D* module. Synapsis between module *Y* and module *K*, and within module *K* are plastic, subject to plasticity rule defined in eq (23), which is applied at all times and is not dependent on reward. Synapsis between module *K* and module *D* are plastic, subject to plasticity rule defined in eq (24), which depends on reward. (**b)** Serial reversal protocol for training of the network depicted in (a). Stimuli are presented for 25 ms, and the motor response to be executed is chosen at *t_decision_
* =15 ms from cue onset. Plasticity between *K* and *D* neurons is applied only if there was reward and within a window spanning from *t_decision_
* to the end of cue presentation.

Conceptually, learning is achieved in two steps. In the first place, neurons in module *K* need to form subpopulations that respond differently to each cue at time *t*, given the past contingency up to the cue presentation at trial *T* −1. This is achieved by plasticity rule described by eq. (23), provided that the system has enough memory so that events in trial *T* −1 have an impact at trial *T*. Next, neurons in module *D* need to read the firing of module *K*, mapping each contingency coded in module *K* to the correct response. This is achieved thanks to the learning rule described by eq. (24), which is proved to reduce the distance between module *D* firing probabilities *p*(*d* | *r*
_1_) and *p*(*d*), leading in turn to an increase in *p*(*r*
_1_) (see Rueckert *et al*.[8]).

The model effectively learns to solve a SRT, as can be seen in Fig. 6a. After 10000 trials of training, the model is capable of changing strategies in the trial immediately following rule reversal (Fig. 6b). The dynamics of synaptic weights along training depends on each kind of connection (Fig. 7).

**Fig. 6.**
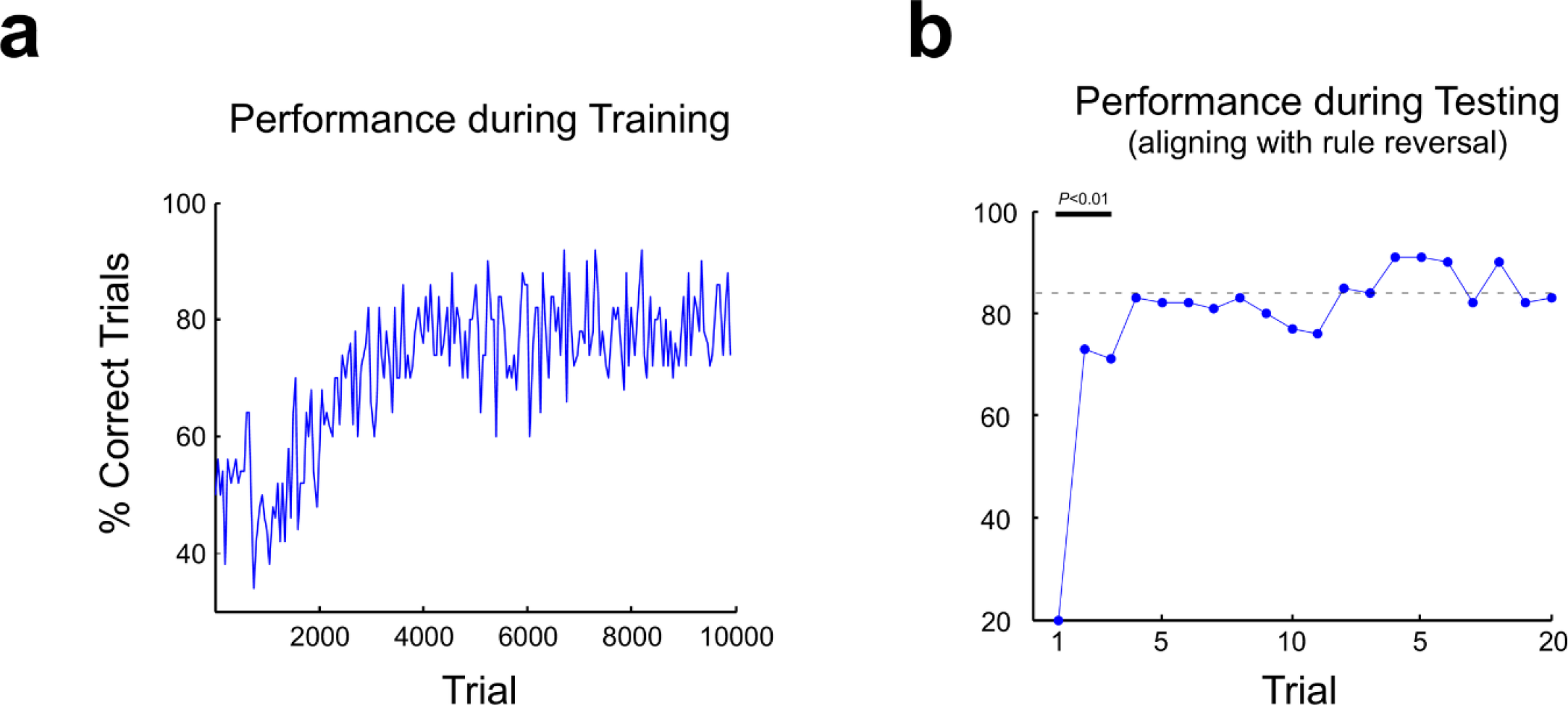
The complex neural network learns to solve the SRT. (**a)** Performance of the model during training, computed as percentage of correct responses in non-overlapping windows of 50 trials. Reversals during training occurred every 15-20 trials. (**b)** The trained model was tested without further plasticity in 2000 trials, with reversals every 20 trials, and performance was computed for each trial, aligning from the trial where the reversal took place. Performance is low immediately after reversal, but improves quickly. Dashed line stands for average performance between trials 4 and 20 from reversal. Performance in the first three trials is significantly lower than average (binomial test, *P*<0.01).

**Fig. 7.**
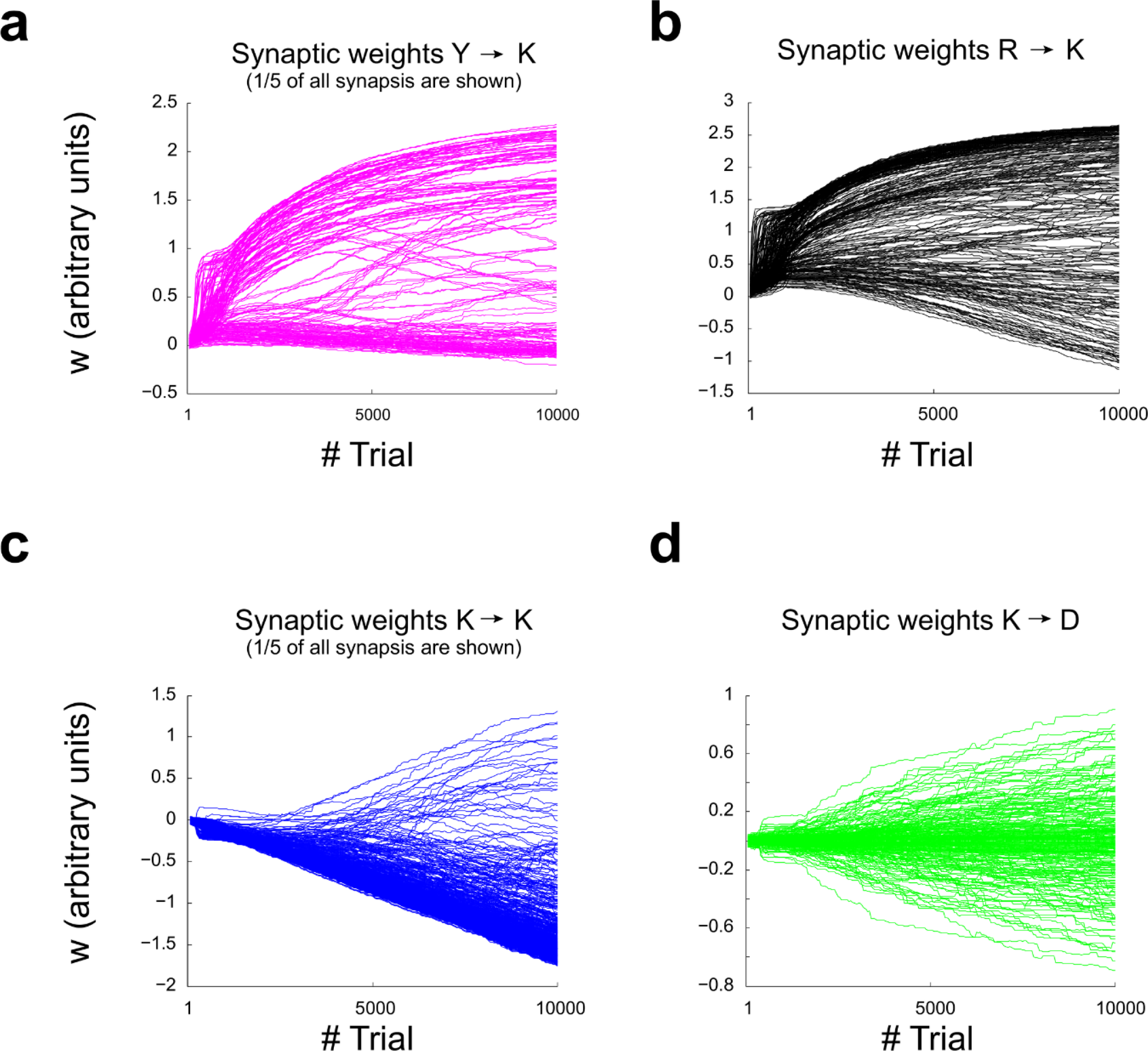
Evolution of synaptic weights of a complex network along training. (**a-b)** Weight distribution for *Y* → *K* and *R* → *K* connections are bimodal, with large values appearing early during training. (**c)** Synaptic weights between *K* neurons follow a strongly skewed distribution, with many small values and a long heavy tail of relatively few large values that evolve slowly during training. (**d)** Connections between module *K* and module *D* follows a symmetric distribution around zero, consistent with the function of module *D* of filtering *K* inputs.

After learning, neurons in module *K* fire in sequences (Fig. 8) which presumably contain the information employed by module *D* to choose the right response. We studied the firing profile of the *K* module by computing the probability of firing of each *K* neuron during *t_decision_
* (Fig. 9a). It can be seen that each one of the 16 possible contingencies has a firing profile that is almost unique. Some contingencies are codified by one single neuron each (for example, contingency 15), while other contingencies are codified by a set of neurons that fire more evenly (for example, contingency 14). This can be seen more clearly by computing a Similarity Index (SI) for firing profiles between all contingency pairs (Fig. 9b). Most pairs have a small SI, and many contingencies are coded by unique sets of neurons. Therefore, the firing state of the *K* module together with the response executed conform a set of states that can be separated in two disjoint subsets when conditioned to reward, which allows the *D* module to map each firing state in module *K* to the correct motor response by means of plasticity rule described in eq. (12).

**Fig. 8.**
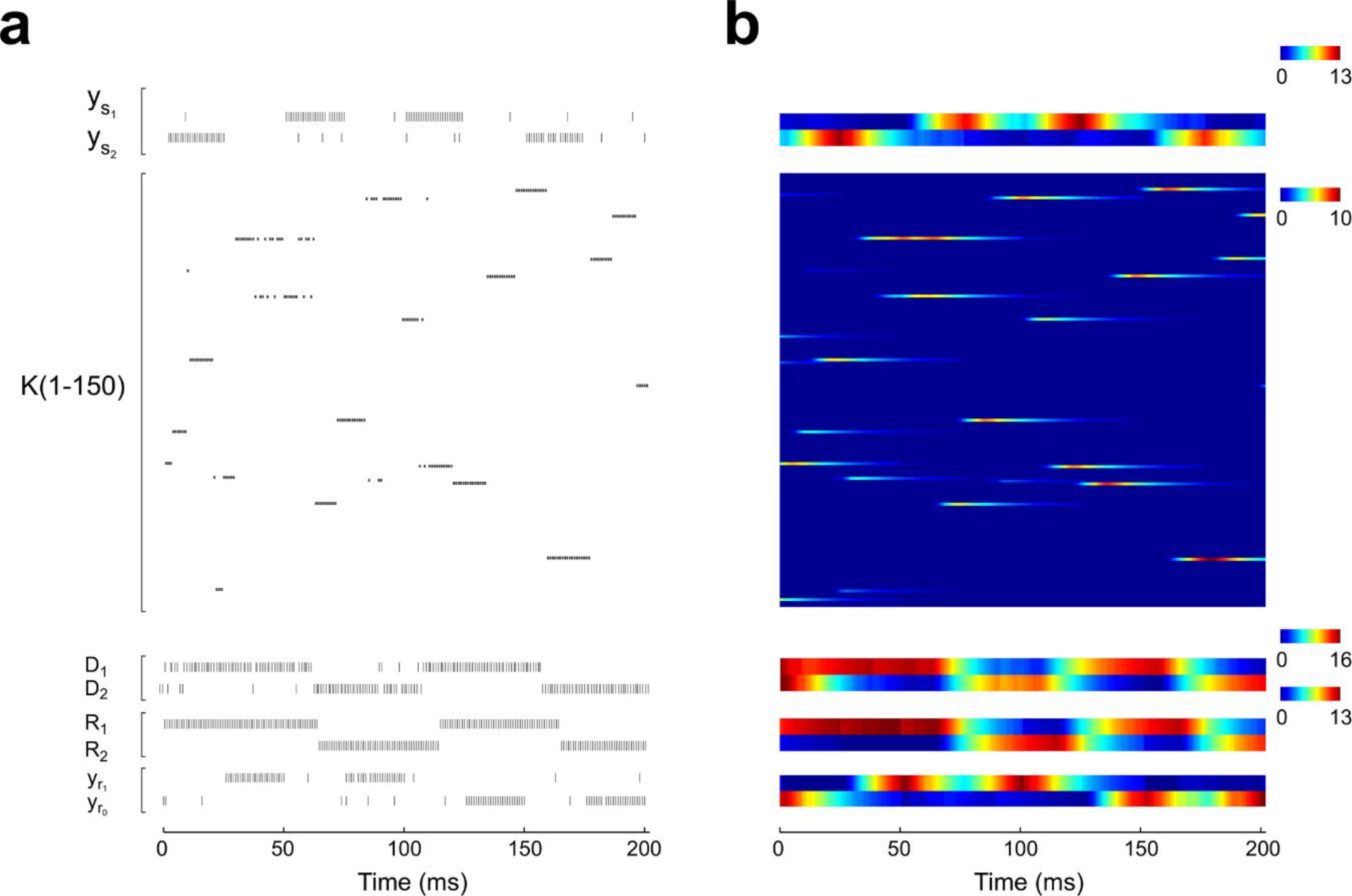
Emergence of sequential firing in the *K* module. Spiking activity (**a)** and corresponding postsynaptic potentials time courses (**b)** of the complex network during 4 consecutive trials of the SRT after achieving high performance. Neurons in the *K* module fire in sequences of sustained bursts of activity. Postsynaptic potentials make each spike has an influence tens of milliseconds after their emission, allowing to link the activity across different stimuli presentations. Note that neurons in the *D* module change their activity after stimulus onset and short before *t_decision_
*. Rule *L*
_2_ was current along the four trials. Colour bars are in arbitrary units.

It is interesting to note that only half of the 16 contingencies are possible within blocks of trials under rule *L*
_1_, being the other half only possible within the block of trials under rule *L*
_2_ This implies that learning the contingencies could be subjected to a problem of catastrophic forgetting. However, this was seldom the case as can be seen from Fig. 9, at least for the protocol of 15-20 trials for each rule. To further explore this issue, we trained networks during 10000 trials under protocols with blocks of crescent number of trials with the same rule, and computed the SI and average performance (Fig. 10a-b). Performance dropt as quickly as the SI values went up, as trials per block were increased, reaching a plateau for the longest blocks. However, it is worth noting that high performance (69%) is still attainable for blocks of 320 trials, showing that the model has a remarkable resilience to catastrophic forgetting of information regarding contingencies.

**Fig. 9.**
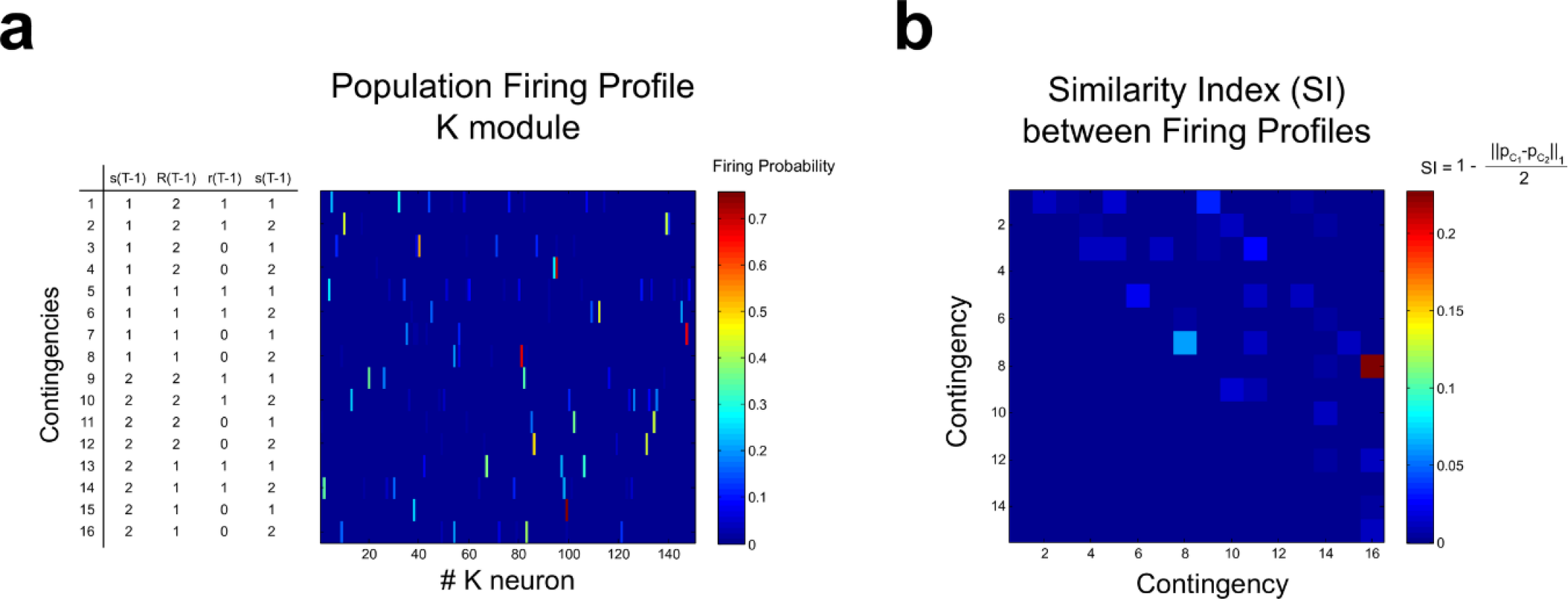
Population coding of stimuli contingencies in module *K*. (**a)** The estimated firing probability of each neuron in module *K* computed at *t_decision_
*, for each one of the 16 possible contingencies. Each row in the heat map represents the population firing profile *p_C_
* for a given contingency *C*. It can be seen that firing profiles do not show significant overlapping. (**b)** Similarity index (SI) between pairs of contingencies firing profiles, which is inversely proportional to the 1-norm between firing profiles, and normalized to the interval between 0 (no similarity) and 1 (total similarity). In general the SI values are low. The highest SI was equal to 0.23, between contingencies 8 and 16, which only differ in their *s*(*T* −1). The second highest SI value was equal to 0.06, computed between contingencies 7 and 8, which only differ in *s*(*T*). There was a tendency for SI values to be high for pairs of contingencies that share the same *s*(*T* −1) or *s*(*T*).

**Fig. 10.**
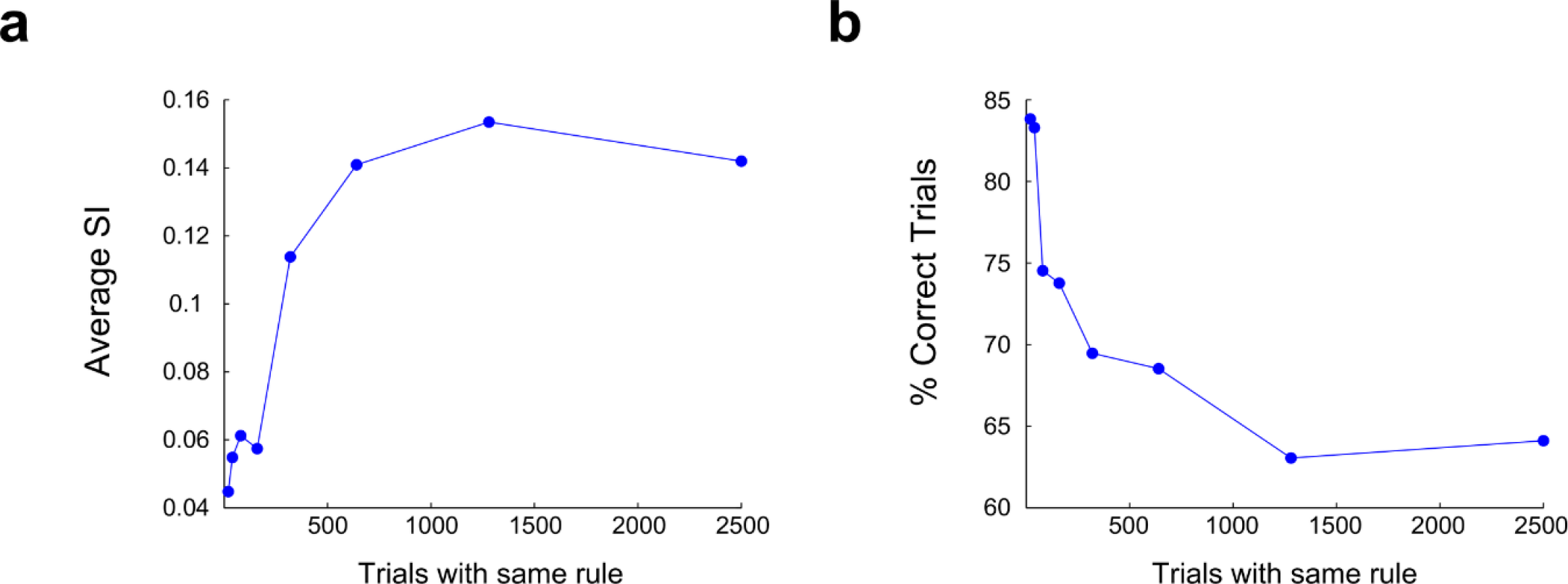
Effect of trials per block on model performance. Networks were trained in the SRT during 10000 trials, and average SI (**a)** and performance (**b)** were computed in 2000 trials without plasticity. Each point in the plot belongs to one network trained with the number of trials per block specified in the x axis. Average SI values were computed from the SI values between pairs of contingencies with shared *s*(*T*) or *s*(*T* −1), which are the contingencies with the highest SI, as shown in Fig. 9.

To better understand the dynamics of learning, we computed how well neurons in module *K* codified each element of the contingency vector (*s*(*T* −1), *R*(*T* −1), *r*(*T* −1), *s*(*T*)) along training. Every 1000 trials of training we employed the last actualized synaptic weights in a separated simulation of 200 trials without plasticity, and assessed contingency coding by training a tree bagger classifier to classify each of the 16 contingency based on the firing of all *K* neurons during *t_decision_
*. Then, the classifier was used to classify trials sharing one of each of the components of the contingency vector (Fig. 11a). Classification performance (CP) before training was around 50% for each separate element, and around 6% for the whole contingency, matching the CP values expected by chance. After 1000 trials of training, the CP of *s*(*T*), *R*(*T* −1) and *r*(*T* −1) were almost 100%. The response stimulus is the only stimulus that lasts 50 ms, and is only changed after *t_decision_
*, meaning that its coding demands the least memory and thus is expected to be the easiest to code, along with *s*(*T*). Coding of reward stimulus *r*(*T* −1) demands more memory from the system, but nevertheless is coded with similar proficiency to that of *s*(*T*) and *R*(*T*). On the other hand, the CP of *s*(*T* −1) grows following a sigmoid-shaped function that resembles the temporal dynamics of the synaptic weights within the *K* module. Within the contingency vector, *s*(*T* −1) is the first stimulus to be presented, and presumably the one having the strongest memory requirements. Moreover, it is followed by the *r*(*T* −1) stimulus, which could act as an interferent. The coding dynamics of *s*(*T* −1) is almost identical to the coding dynamics of the entire contingency vector, and also grows similar to the growth in behavioural performance (Fig. 6a), suggesting that coding *s*(*T* −1) is the bottleneck for contingency coding, and presumably for behavioural learning.

**Fig. 11.**
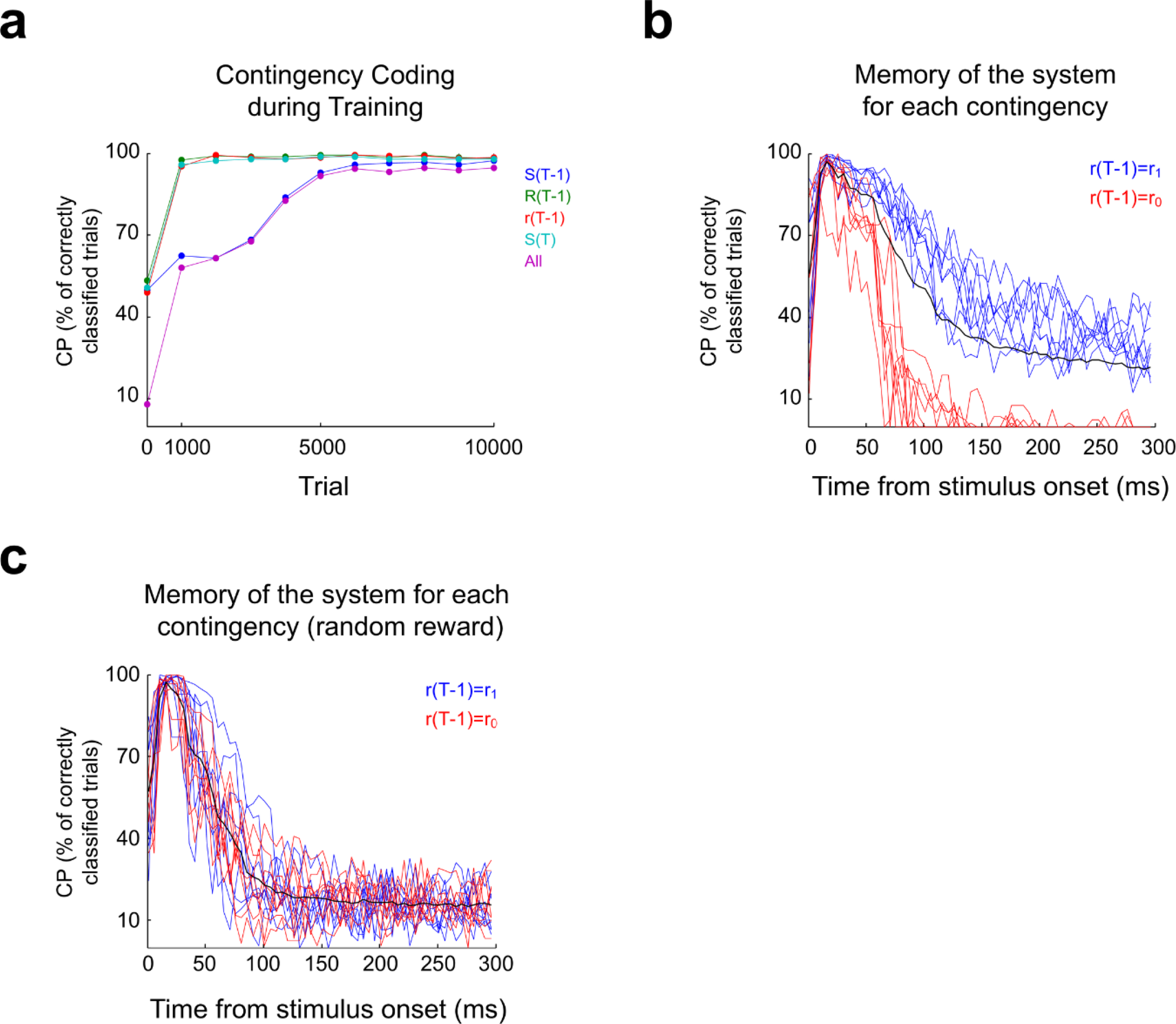
Contingency coding and memory after training. (**a)** The information conveyed by the *K* module about the contingencies was estimated by employing tree-bagger classifiers trained on the *K* module firing profile to classify trials according to their membership to a given group of contingencies that share some specific element, depicted in the legend. Probe simulations were run before beginning training (Trial = 0) and then every 1000 trials. Firing profiles where computed at *t_decision_
*. Information about *s*(*T* −1) takes more training to be acquired, acting as a bottleneck for the coding of the whole contingency. (**b)** Memory about the occurrence of each contingency was estimated by assessing the classification performance of a Naive Bayes classifier trained to correctly classify the 16 contingencies based on the *K* module firing profile computed from *t* = 0 of trial *T*, to the end of trial *T* + 5 (being *T* the trial when *s*(*T*) of the target contingency was presented). The CP value picks around *t_decision_
* as expected since contingency may change after that time. For contingencies which involved the *r*
_1_ stimulus, information is retained above chance levels long after the time of decision. On the contrary, information about contingencies involving *r*
_0_ was retained for a shorter period, suggesting that information retention is proportional to the frequency of occurrence of the contingency. (**c)** When reward is delivered at random, differences in information retention between contingencies involving *r*
_1_ and *r*
_0_ disappears.

Results in Fig. 11a show that module *K* has enough memory to retain information for at least 50 ms. To further explore the memory capacity of the system, we tested the model that learned the SRT by simulating 2000 trials without plasticity. Trials were sorted according to their membership to each contingency and a Naive Bayes classifier was trained to classify trials according to their membership to a given contingency, based on the activity of the *K* neuron population at time points ranging from the start of *s*(*T*) to the end of *s*(*T* + 5) presentation (300 ms of consecutive activity). The CP was assessed for each contingency separately, and for the set of 16 contingencies (global performance), (Fig. 11b). Global performance starts around 50% at *t* = 0 ms, which means that the *s*(*T* −1), *R*(*T* −1) and *r*(*T* −1) components were already codified at trial initiation; uncertainty remained regarding *s*(*T*), which is expected since this stimulus had not been presented at *t* = 0 ms. Global performance picked rapidly, reaching its maximum of 97% at *t* =15 ms. At this time, the response is actualized and thus can differ from the response in the contingency being analysed, explaining that the maximum global performance is found at *t_decision_
*.

For good performance, information about the previous trial must be retained until *t_decision_
*. We can see that the memory of the system far exceeds this minimum requirement, with a CP of 22% at *t* =300 ms. Notably, CP values per contingency are clustered in two well defined groups that differ in how quick classification performance drops. The group of contingencies for which the system has shorter memory (CP drops fast) is composed of contingencies where *r*(*T* −1) = *r*
_0_ (red curves), while memory is longer for rewarded contingencies. It is important to note that *r*
_0_ is less and less presented as learning progresses, leading to an underrepresentation of contingencies containing *r*(*T* −1) = *r*
_0_. This suggests that the number of times a given contingency was presented during training defines for how long the system retains information about that contingency. To test this hypothesis, we performed a new training in which both *r*
_1_and *r*
_0_ have equal chances of been presented regardless of the chosen motor response. In this case, the CP of all contingencies followed a similar temporal course, as expected (Fig. 11c).

## Discussion

In this work we have studied under what conditions a biologically plausible neural network is capable of solving a serial reversal task. The distinctive feature of this paradigm is that each stimulus/response pairing is eventually reinforced, since correct responses depend on the current rule. Thus, the sole information about the perceived stimulus and executed response collected at any single point in time is not sufficient to solve the task. This problem is reminiscent of the problem of catastrophic forgetting, also called the stability/plasticity dilemma, which is usually stated as the difficulty that many neural networks models have in acquiring new information without erasing old information [2,9]. Catastrophic forgetting studies usually focus on paradigms where a set of stimulus response pairings must be learned sequentially. Thus, the difficulty of the task strives in the distributed representations of stimuli in the neural network, where the same set of synaptic weights are modified each time a new pairing is presented. It has been shown that forgetting can be alleviated in models that incorporates different levels of plasticity, i.e. mataplasticity [10,11]. Moreover, previously acquired information can be preserved in the correlated firings of the neural population [12]. Thus, it might be reasonable to think that similar mechanisms could be at work in a related behavioural paradigm like the SRT. However, the results presented in this work show that no plasticity rule or neural activation function is sufficient to guarantee good performance in the SRT, without contradicting the hypothesis of functionality by learning. In particular, we showed that the SRT cannot be learned by any network in which the same neural population integrates stimuli information, and at the same time defines the motor response through non-plastic connections.

It is assumed that learning occurs through neural mechanisms that drive the network to a configuration that solves the task. A prerequisite for learning is that the probability of *sequences* of stimuli and responses must be different when conditioned to reward than when conditioned to no reward; the non-fulfilment of this prerequisite means that reward delivery is not dependent on behaviour and there is nothing to be learned. Then, the network must achieve two properties: to differentially code in its states the sets of rewarded and non-rewarded sequences of stimulus/response pairings, and to map network states to the correct motor response. It is important to note that this last property (mapping) is only attainable after the first property (coding) is achieved. In the simple network, once coding is achieved the mapping is completely defined, since motor responses are pre-defined based on the activity of the integration/decision module *K*. But adequate mapping requires appropriate coding as a prerequisite, implying that simple networks will achieve mappings that allow high performance only by chance, which although not impossible, since we considered stochastic networks, is something that can hardly be regarded as learning. Moreover, the probability of finding a solution in this way would be very low, since the solution trajectories are only a small subset of all possible trajectories. For example, the module *K* ruled by equations (18-24) is capable of coding the 16 (*s*(*T* −1), *r*(*T* −1), *R*(*T* −1), *s*(*T*)) sequences. Let’s considered 16 *K* neurons, half of them leading to *R*
_1_ and the other half leading to *R*
_2_. We may assume that each neuron will code one of the 16 possible contingencies at random, since initial conditions were randomly chosen. Then, there are 8! x 8! out of 16! possible assignments between *K_i_
* neurons and contingencies *C _j_
* that lead to 100% of correct responses. This mean that, by choosing an initial random condition, this *K* module will exhibit 100% performance with a probability of 7.8x10^-5^.

Building from the restrictions exhibited by the simple network scheme, we proposed a neural network model capable of solving the SRT, which relies in assigning the functions of contingency coding and response selection to different neural populations (integration module *K* and decision module *D* in Fig. 5a). In this way, all the information required to solve the SRT (i.e. the coding of the (cue,response,reward) contingencies) is firstly acquired in module *K*, and then the module *D adapts* its response through reinforcement learning in order to maximize reward.

It is interesting to note that, although the separation of functions achieved in the complex model allows to untie the problem generated by the reversal paradigm, the coding of the contingencies themselves implies a possible problem of catastrophic forgetting, because it is the same set of synaptic weights that is required to change to learn contingencies which are presented in a sequential schedule. Nevertheless, the soft winner-take-all network implemented as module *K* showed a remarkable resilience to forgetting. Although information of contingencies within a block of trials with the same rule could persist long enough into the other block, this is not likely, since the memory of the system declines considerably after 6 trials (Fig. 11b). A possible explanation for the resilience to forgetting could be found by noticing that the distribution of synaptic weights attained among neurons in module *K* is sparse, a fact that could lead to decreasing the chances of interfering representations [13].

The impossibility result shown here has special meaning for brain regions typically related to decision making like the prefrontal cortex (PFC). The PFC is key to several high level cognitive process such as behavioural plasticity [14], working memory [15–17], rule learning [18] and decision making [19]. Its function is often described as of integration of information and response selection. For example in Mante *et al*. [20] a target cue informs in each trial which sensory dimension is relevant for obtaining reward, and the behavioural and electrophysiological data is explained in terms of a single neural network population that integrates both relevant and irrelevant sensory information and choose the right motor response, given suitable synaptic weights that should be attained through learning. Other commonly employed task like delayed matching-to-sample can also be solved by a single neural population model as the one depicted in Fig. 1b. Nevertheless, the PFC was also shown to be necessary for the SRT [21]. This suggests that in order to understand the PFC role in the SRT, and in IRTs in general, it is important to take into account the subpopulations within the PFC, from the level of micro circuits to the interconnections between PFC subregions. Specially, it would be of interest to characterize subpopulations of neurons according to their afferent and efferent projections and in relation to their firing profile. It could be expected that the PFC neurons could be sorted in populations of coding neurons, that code complex contexts and stimuli histories, and decision neurons, that integrate contingency information from the coding population and projects to motor structures like the basal ganglia, or the motor cortices.

Synaptic plasticity in the model depicted in Fig. 5a fulfils two different functions. In module *K*, plasticity allows the system to classify stimuli contingencies. The modulation of plasticity by reward would make no difference there, since all contingencies are equally rewarded, at least during the beginning of learning. Evidence of sustained plasticity have been found experimentally, in the form of the continuous formation and erasure of synaptic spines in cortex, which occurs even in the absence of any obvious reward [22]. On the other hand, plasticity between module *K* and module *D* has the function of allowing the *D* module to read the firing of *K* neurons that carries contingency information, and to map it with the correct response. In this case a reward-modulated form of synaptic plasticity is essential, and related experimental evidence can be found in the known effects that the neuromodulator dopamine (DA) has on synaptic plasticity in brain regions like the cerebral cortex [23], hippocampus [24] and striatum [25], and in the fact that DA neurons code reward and reward-predicting cues [26,27]. This fundamental difference in plasticity modes in the model suggests that experimental approaches to understand neural computation should focus on searching for subpopulations based in their synaptic plasticity profile, dissecting populations of neurons according to how sensitive their synaptic changes are to neuromodulators related to reward. Understanding the relationship between connectivity, firing profile, and reward and non-reward modulated plasticity could help to discover the building blocks of neural computation.

In brief, the study of a well-known task as the SRT allowed to gain new insights into the computational limits of an important set of biological neural networks that are commonly considered as models of learning and decision-making, and to give new theoretical support to the experimental exploration of the anatomy and function of neural circuits. Future work should focus on the rules of connectivity that allows greater memory for coding more complex contingencies, and in the kind of algorithms that can be learned by combining different circuit motifs with reward and non-reward modulated plasticity.

## Methods

### Proofs for the impossibility of simple neural networks to learn to solve the SRT

Achieving high performance in the SRT implies that the network responds to stimuli according to the transitions depicted in Fig. 2. The behaviour of the network will be inherently stochastic, since it is required to respond to stimuli that are themselves stochastic. However, given the state of the network at time *t*, the transition probability for the correct response is expected to be close to one, with all other responses having transition probabilities close to zero. Without the stochasticity of the stimuli, the network would follow a deterministic limit cycle, in which *n*(*t*) = *n*(*t* + *m*), being *m* the length of the cycle. In this manner, we say that the transition probability matrix is a deterministic probability matrix, and that the network follows a stochastic limit cycle, where the stochastic component of the behaviour is given by external factors that do not depend on the activity of the network.

With this concepts in mind, we will prove that the reduced neural network cannot learn to solve the SRT by showing that, given any excitation function *f* and plasticity function *g*, the network either does not converge, or it converges to one of many possible stochastic limit cycles, where only a small subset of these limit cycles allow high performance in the SRT.

First, we will study the convergence properties of the reduced neural network, assuming that external stimuli are not stochastic. We build on the mathematical framework of decision systems as presented in [28]. There, a decision system is defined, which is composed of a state space *X*, a decision space *D*, and transition probabilities *p_i_
* (*x*):= *p*(*i* | *x*) and *P_i_
* (*x*, *A*):= *p*(*x* ∈ *A* | *x*,*i*), where *x* ∈ *X*, *i* ∈ *D*, and *A* is any element of the sigma algebra on *X*. At each time *t*, a decision *i* is taken given *x*(*t*) and *p_i_
* (*x*), obtaining *i*(*t* +1). Then, *x*(*t* +1) is obtained, conditioned to *i*(*t* +1) and *x*(*t*) through *P_i_
* (*x*(*t*), *x*(*t* +1)).

The evolution of a stochastic spiking network can be represented within this framework by the following representation:

*D* = {(*z^i^
*, *z ^j^
*)}, the set of all possible pairs of firing network states the network can assume. Vectors *z^i^
* is the ith vector of the set of all possible firing state vectors *z*

*X* = {(*w*, *z*, *rew*)}, the set of all possible combinations of whole system synaptic weights configurations (*w*), networks firing states (*z*) and reward function *rew*.

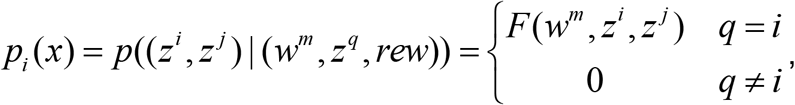

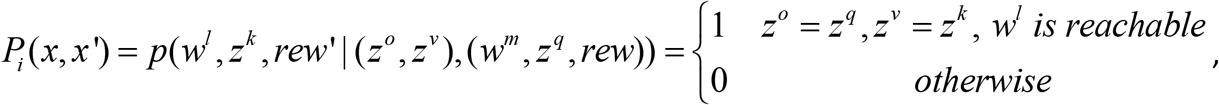

where *w^i^
* is the ith vector of the set of all possible synaptic weight configurations for the whole network. By *reachable* we mean that *w^l^
* is the whole synaptic configuration that is obtained when applying plasticity function *g* after transition from *z°* to *z^v^
*, having *rew* the value corresponding to that trial given *z^v^
*, *s* and *L*.

Theorem 1 in [28] shows that a decision system converges with probability 1 to a limit cycle if and only if for each state *x* there is a decision *i* such that:

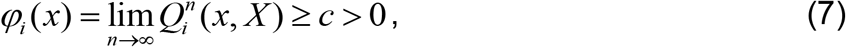

where *c* is a constant and 
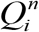
 is defined inductively as 
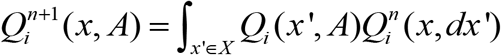
, with 
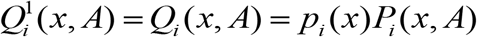

Intuitively, condition (7) is fulfilled only when the probability of transitioning from firing state *z^i^
* to *z ^j^
* infinitely often does not vanish, which happens only if the probability converges to 1.

In the case of a reduced network

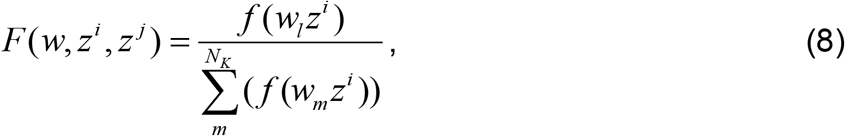

where *l* is the *K* neuron that is active in state *j*. The function *f* is any function with the condition that is strictly increasing with *w_i,j_
* ∈ ℝ.

It is the fact that synaptic weights change deterministically the reason why *p_i_
*(*x,x’*) is either 1 or 0. This allows us to simplify condition (7) to:

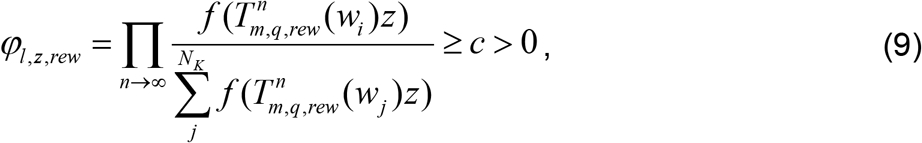

where *l* is the active neuron in the destination state *j*, *z* is the source state 
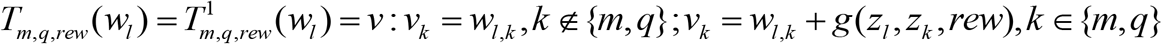
, and 
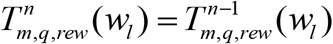
 The transformation *T* takes the vector of synaptic weights of inputs to neuron *l* and applies a synaptic change according to plasticity function *g* (eq. 2) to the weights corresponding to presynaptic neurons *q* and *m* (one for a neuron *Y*, the other for a neuron *K*) that are active, i.e. *z_m_
* = *z_l_
* = 1.

Equation (9) holds only if log*φ_l_
*, _
*z*,*rew*
_ converges, which is an infinite sum of logarithms. In turn, the sum converges if the application of *T* leads to an increase in the transition probability. Since *f* is strictly increasing with *w_i_
*, _
*j*
_, eq. (9) holds if *δ*
_1,1,*rew*
_ > 0. In other words, the network will converge to a limit cycle if for each pair of active neurons *Y* and *K* there is a transition to a neuron *n_l_
* such that, if the transition is repeated infinite times, the probability of the transition increases, something that occurs if the pre/post activation leads to potentiation of the synapse, i.e. Hebbian plasticity.

As stated before, a neural network that must learn to solve the SRT will not reach a limit cycle since it is bonded to follow stimuli that are stochastic. However, the evolution of the network can be segmented in transitions that eventually reach probability 1. Namely, for states define as (*s*, *n_i_
*, *L*) and (*r*, *n_j_
*, *L*), we can consider transitions conditioned to given *s* and *L*, i.e. the external stochastic factors which are independent of the network behaviour. For example, the transition between a given source state (*s*, *n_i_
*, *L*) and destination states (*r*, *n_j_
*, *L*) for any neuron *n _j_
*, can be considered a decision system. Then, if condition (9) is fulfilled, any of these decision systems will converge to a “limit cycle” in which only one destination state (*r*, *n_j_
*, *L*) is chosen. The same holds for transitions between source state (*r*, *n_i_
*, *L*) and destination states (*s*, *n_j_
*, *L* ’). It is important to note that a neural network that solves the SRT needs to converge to a unique decision even for incorrect trials, i.e. for source states (*r*
_0_, *n*, *L*). This means that *g*(*n_i_
*, *n_j_
*, *rew*) > 0 for any *rew*∈{1, 0}.

Any pair of source and destination states can became the limiting transition, the probability of this happening depending on the initial transition probability, which depends on the initial synaptic weights. In particular, for networks in which eq. (9) holds, any limiting transition is attracting since the transition probability rises with probability equal to itself. The SRT is solved with high performance for only a small subset of all the possible limiting transitions. Therefore, a simple network which is initialized with random synaptic weights will reach a synaptic configuration that solves the SRT with very low probability. In particular, the probability of reaching the solution will be high only if the initial transition probabilities are close to the solution probabilities.

### A more general definition for the simple neural network

The reduced neural network can be extended to a more general definition of simple network, with arbitrary number of neurons and for which the impossibility result holds. In this case, the networks dynamics develops in discrete time steps of 1 ms each. The SRT is structured in trials composed of cue stimulus presentation followed by a reward stimulus presentation, each one lasting *t_stimulus_
* in ms. The response is observed in the interval [*t_cue offset_
* − Δ*t_response_
*, *t_cue offset_
*]. The sequence of firing states of module *K* during this time interval univocally defines the behavioural response *R*.

We will consider a simple neural network composed of *N_Y_
* neurons in module *Y* and *N_K_
* neurons in module *K*. The firing state of the ith neuron in module *Y* will be represented by the variable *y_i_
*, the ith element of vector *y*. The firing state of the ith neuron in module *K* will represented by the variable *n_i_
*, the ith element of vector *n*. Neurons in module *Y* fire independently of each other, conditioned to the stimulus presented:

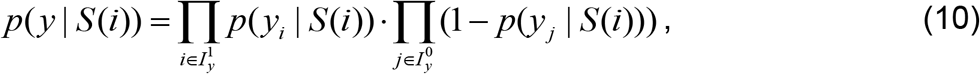

where 
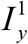
 is the set of indexes of *Y* neurons that are active in vector *y*, and 
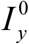
 is the set of indexes of *Y* neurons that are inactive in vector *y*.

The postsynaptic potential *PP_i_
*, _
*j*
_ elicited by the train of spikes of neuron *j* onto neuron *i* is defined as the product of the post synaptic potential time course: *x _j_
* and the corresponding synaptic weight

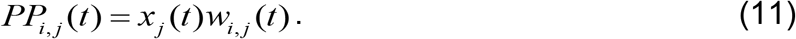

The variable *x _j_
*, the postsynaptic potential time course associated with the spike train of neuron *j*, is defined as:

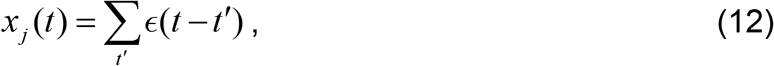

Where ∈ is a kernel function, and *t*′ runs over all the firing times up to time *t* at which the jth neuron of the module fired.

The excitability of neuron *i* in module *K* is defined as:

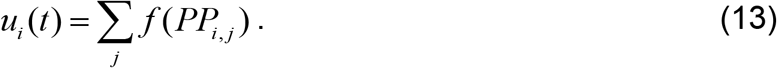

Conversely, its probability of firing is:

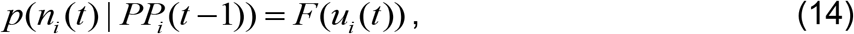

Where *sign*(*f* (*PP_i,j_
*)) = *sign*(*PP_i,j_
*), 
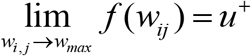
 and 
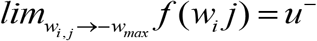
. The function *F* is such that 
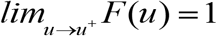
, 
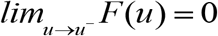
, *F(u)* = 1 for u ≥u^+^ and *F(u)* = 0 for *u* ≤ *u*
^−^. In this way, neuron *i* will fire with probability 1 with the sole firing of a neuron *j*, provided that *w_ij_
* is maximal, and will remain silent with probability 1 if *w_ij_
* is inhibitory (negative) and maximal in absolute value.

Any number of neurons may fire at the same time, and all neurons are conditionally independent of each other given *PP*. Thus, the probability of an activation state *n(t)* whole module *K* is given by:

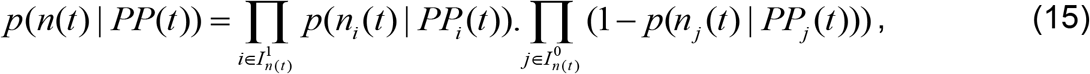

Where 
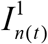
 and 
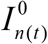
 are sets of indexes of neurons that are respectively active and inactive in *n*(*t*).

Neurons are plastic all the time. Synaptic weight *w_i_
*, _
*j*
_ changes according to the function *g*, defined as:

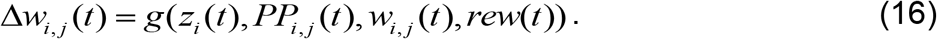

The Δ*w_i,j_
* values depend on *w* in such a way that 
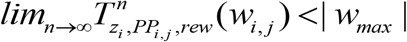
 This assures that synaptic weights remain within reasonable prefixed limits. In this case, the variable *rew* is defined as:

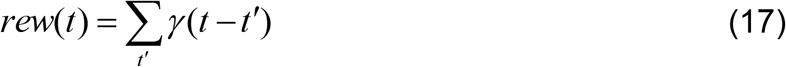

with *γ* a kernel function and *t*′ the onset times of stimulus *r*
_1_.

For a simple neural network defined according to eq. (10) to (17), the impossibility result holds. In particular, since the plasticity rule is deterministic, transitions with probability one will be possible if the corresponding value of Δ*w_i_
*, _
*j*
_ is positive. In this case, all the variability in the network will stem from the stochastic nature of stimuli presentation and rule switching, and from the uncertainty in the coding of stimuli by the sensory module *Y*.

### Implementation of the complex network

In the implementation of the complex network sketched in Fig. 5a, module *Y* was composed of two *Y* neuron for coding each cue stimulus, two *Y* neurons for each reward stimulus, and one neuron *Y* for coding each response. Module *K* was composed of *N_K_
* = 150 neurons. All initial synaptic weights were sampled from a normal distribution of mean=0 and standard deviation= 1/64. There were no self-connections (*w_i_
*,_
*i*
_ = 0).

Each neuron *i* in module *K* has a variable *u_i_
*:

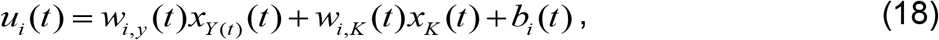

Where *w_iy_
* is a vector containing the synaptic weights for the connection from each neuron in the module *Y* to the ith neuron in module *K*, while *w_iK_
* is an analogous vector for the inputs that neuron *i* receives from the other neurons in module *K*. The vector products *w_i_
*, _
*y*
_ (*t*)*x_Y_
* (*t*) and *w_i_
*,_
*K*
_ (*t*)*x_K_
* (*t*) represent the postsynaptic potentials (*PP*) at time *t* associated with the train of spikes at each afferent synapse from module *Y* and *K*, respectively. The ith element any vector *x* represents the temporal course of the *PP*, which only depends on the spike emission times, and is defined as:

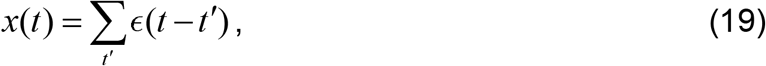

where *t*′ runs over all the firing times up to time *t* at which the ith neuron of the module fired, and ∈ is a double exponential kernel function:

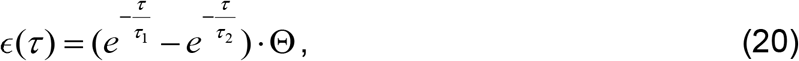

where *τ*
_1_ =2 ms, *τ* _2_ =20 ms, and Θ stands for the Heaviside function. The parameter *b_i_
* controls the excitability of the neuron. This parameter was adjusted at each time *t* following the homeostatic mechanism described in Habenschuss *et al*. [29]:

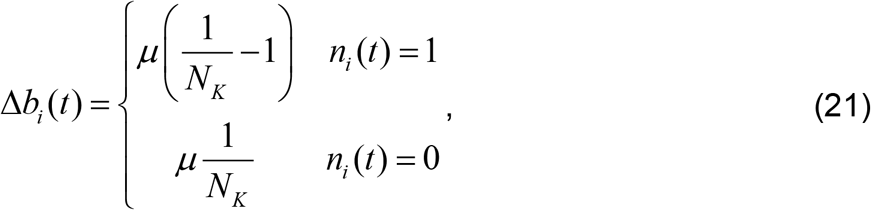

which assures that each neuron in the module fires with equal probability, helping to exploit all neurons in the module, avoiding silent neurons and thus favouring learning. The parameter *μ* was set to 0.1.

The firing probability of neurons in the *Y* module where defined by the stimulus they coded, such that 
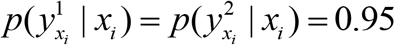
 and 
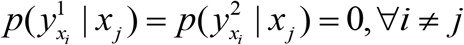
, where 
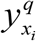
 is the qth *Y* neuron coding stimulus *x_i_
*. The response executed was coded by one *Y* neuron each, such that 
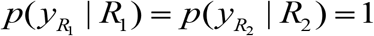
 and 
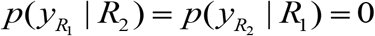

Within module *K*, the firing probability of neuron *i* is defined as:

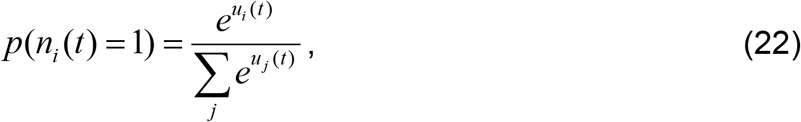

with index *j* going through all neurons in module *K*.

The firing probability of the two neurons in module *D* are defined just as for neurons in module *K*, with the sum in eq.(22) encompassing only the two *D* neurons. Only one neuron in module *K* and *D* fires at each time *t*.

Connections from module *Y* to module *K*, from module *K* to module *D*, and between neurons in module *K* are plastic. The connections from neurons *Y* to neurons *K* and between neurons in module *K* change at each time *t* according to Δ*w_ij_
*:

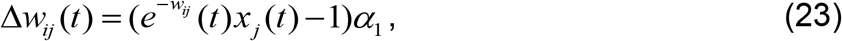

where index *i* refers to the postsynaptic neuron, index *j* to the presynaptic neuron, *x* is the time course of the postsynaptic potential associated with neuron *j*, and *α*
_1_ =5x10^-4^ is a learning constant. This plasticity rule is a kind of STDP rule that leads the model to codify each stimulus by a different population of neurons. Note that the rule does not depend on reward, and weight changes are applied at each time *t*.

Connections from module *K* to module *D* change over time according to:

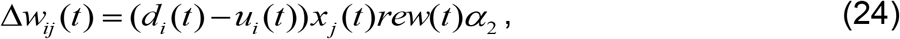

where *d_i_
* stands for the firing state of decision neuron *i*, *u_i_
* is its excitability variable, *x _j_
* the *PP* time course of afferent neuron *j* and *α*
_2_ =8x10^−4^ is a learning constant. The variable *rew* equals 1 only during the decision window and only if the motor response was respectively correct. Otherwise, *rew* = 0. Note that this plasticity rule is conditioned to reward while rule in eq. (23) is not.

### Simulations and analysis

In general, a training session in the SRT consisted of 10000 trials, while a test session consisted of 2000 trials. For the results in Fig. 10, one network per point in the plot was trained during 10000 trials. Each of these training had a specific (fixed) number of trials per block with the same rule, starting from 20 trials per block and increasing the number by factors of powers of 2.

The similarity index employed in Fig. 9b was defined as:

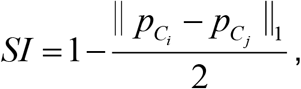

where *p_C_
* is a vector in which the ith element is the estimated probability of firing of neuron *i* conditioned to contingency *C*x. The SI adopts values from 0 (when firing probabilities under both contingencies are equal for each neuron) to 1 (when every neuron fire with probability 1 under one contingency, and with probability 0 under the other contingency.

We employed classifiers to obtain a measure of the information conveyed by the neuron population of module *K* about contingencies. Specifically, for the result shown in Fig. 11a we employed the *TreeBagger* function in Matlab R2009b to train 50 trees, to match the firing of the *K* module at *t_decision_
* with their corresponding contingency. The classification performance was obtained as CP=100x(1-err), where err is the out-of-bag misclassification probability, obtained through the *oobError* function. For the result shown in Fig. 11b we employed the *NaiveBayes* function. We trained 100 classifiers onto 80% of each training set and tested performance in the 20% remaining. The CP in this case was the average performance of the 100 classifiers in the test set, expressed as percentage. The results shown hold regardless of which classifier was employed.

## Acknowledgments

We thank Sergio Lew for helpful comments.

